## Supplementary figures and images for "Sequence features of retrotransposons allow for epigenetic variability"

### Figure 1 - figure supplement 1

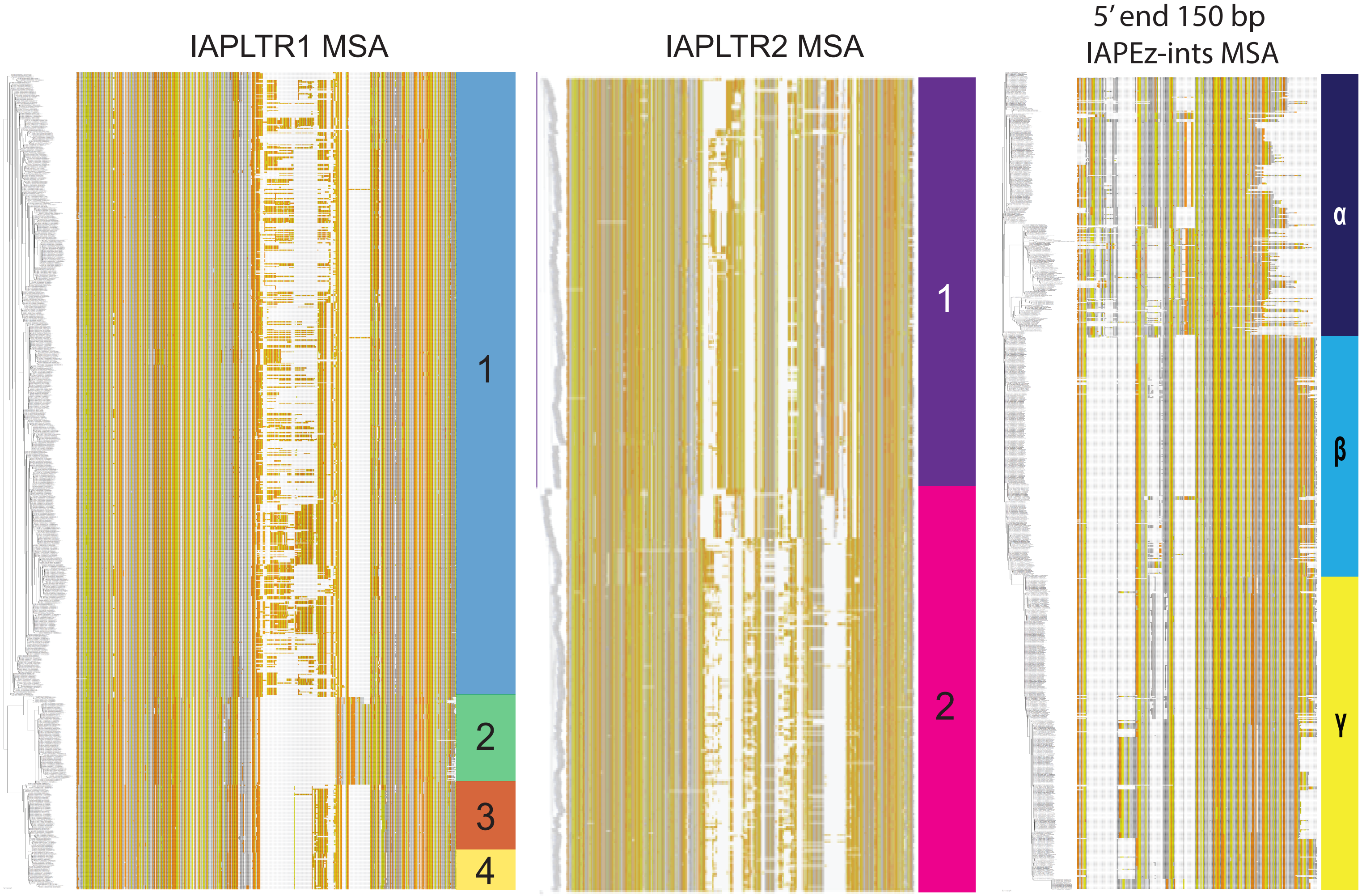

### Figure 1 - figure supplement 2

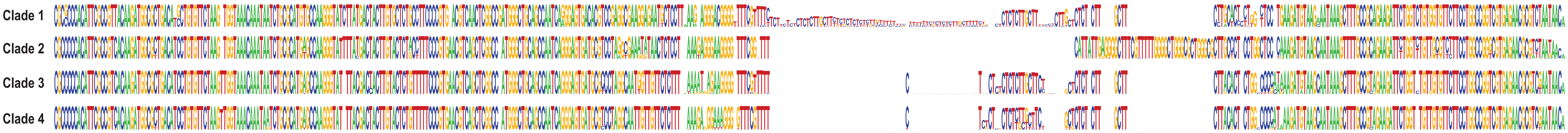

### Figure 1 - figure supplement 3

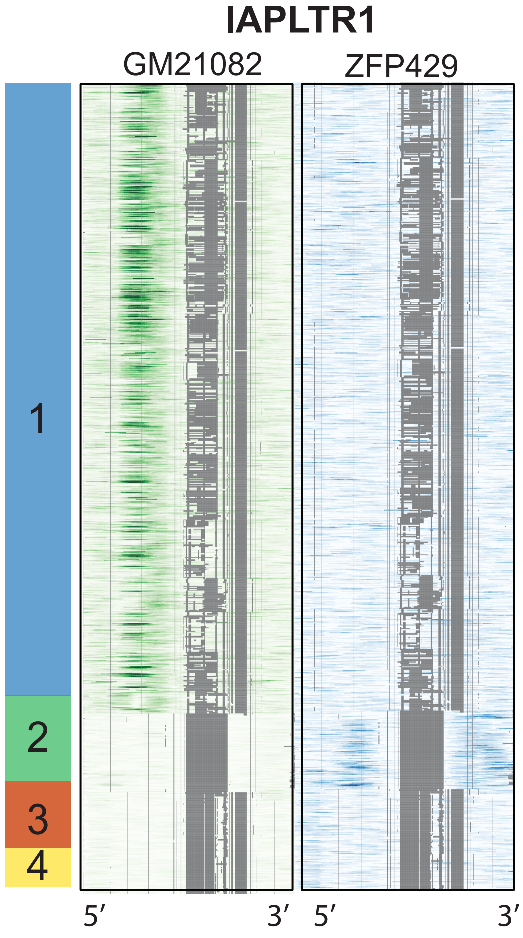

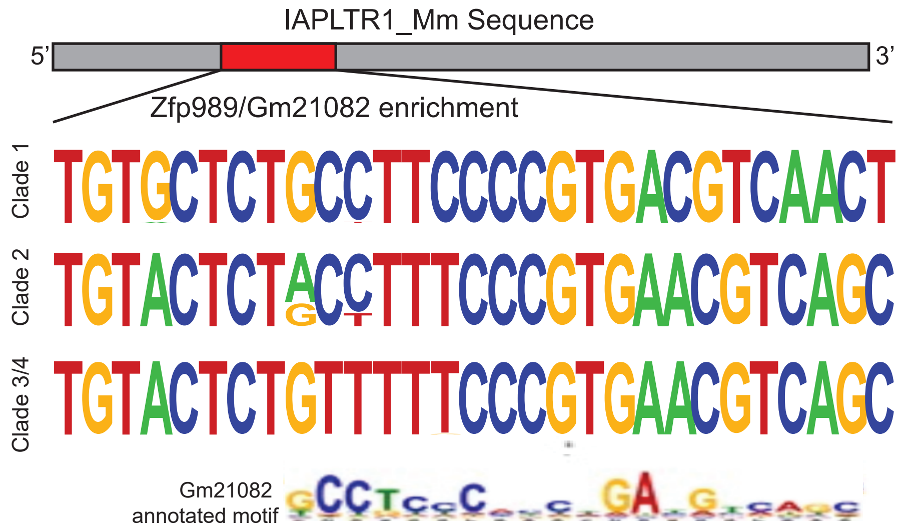

### Figure 1 - figure supplement 4

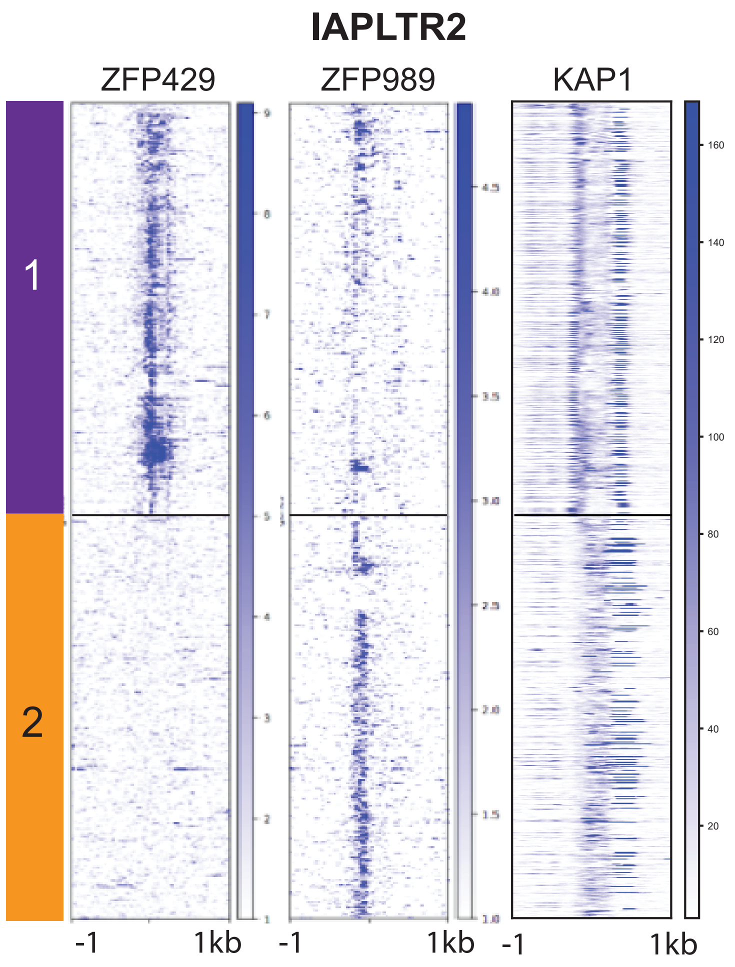

### Figure 1 - figure supplement 5

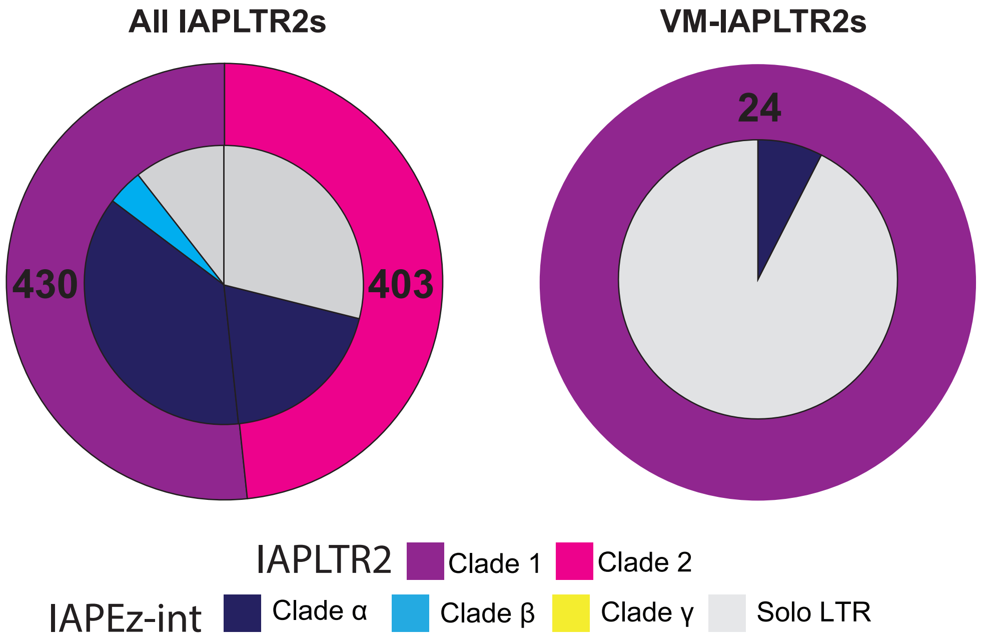

### Figure 1 - figure supplement 6

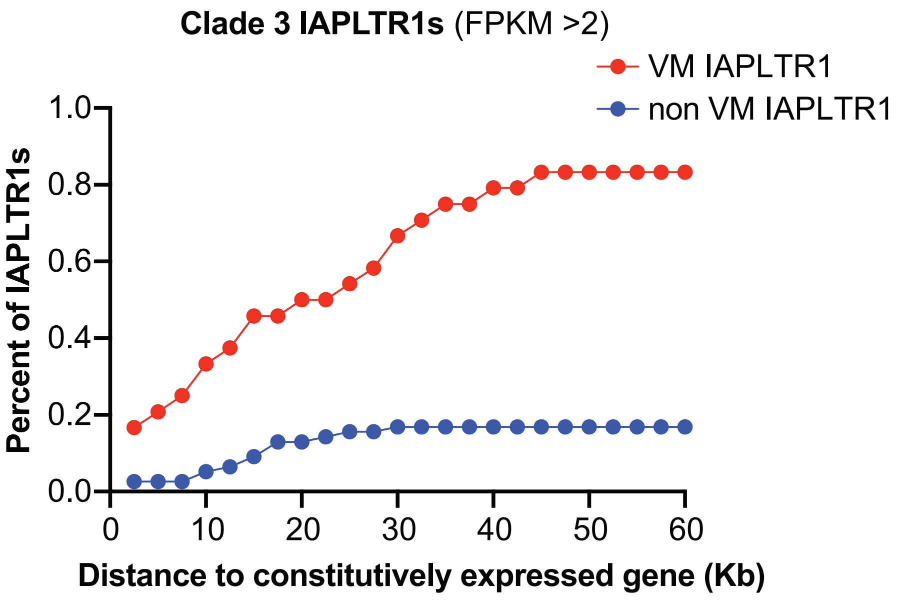

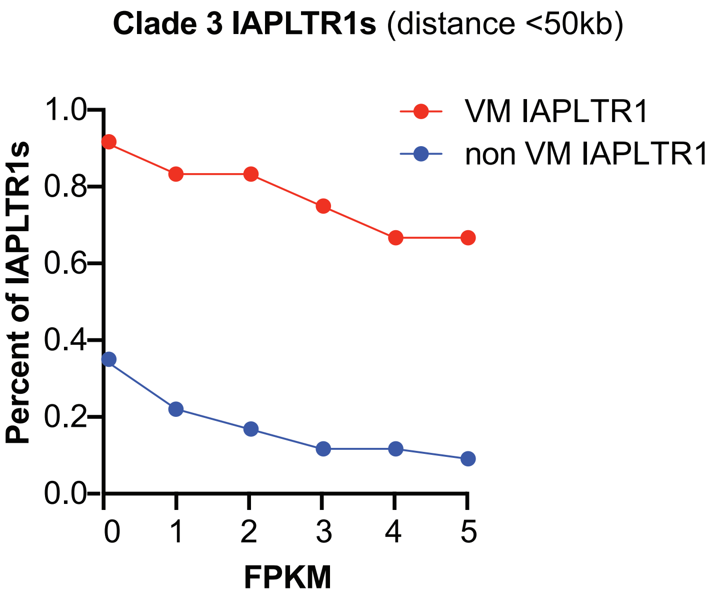

### Figure 1 - figure supplement 7

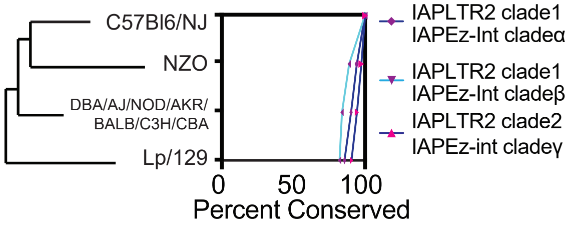

### Figure 1 - figure supplement 8

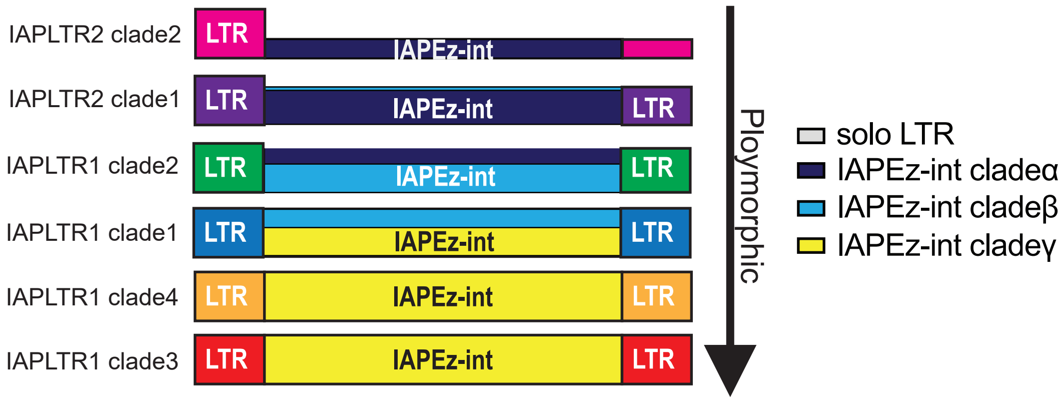

### Figure 2 - figure supplement 1

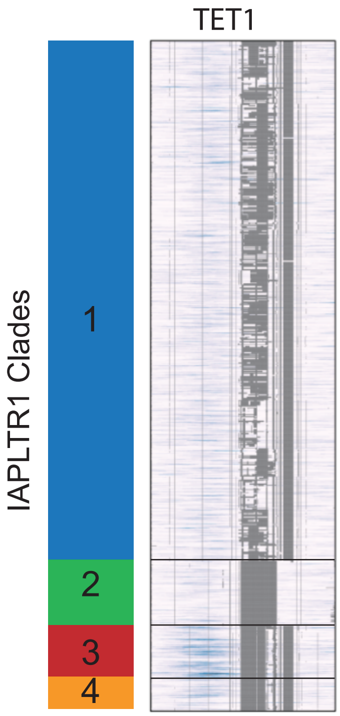

### Figure 2 - figure supplement 2

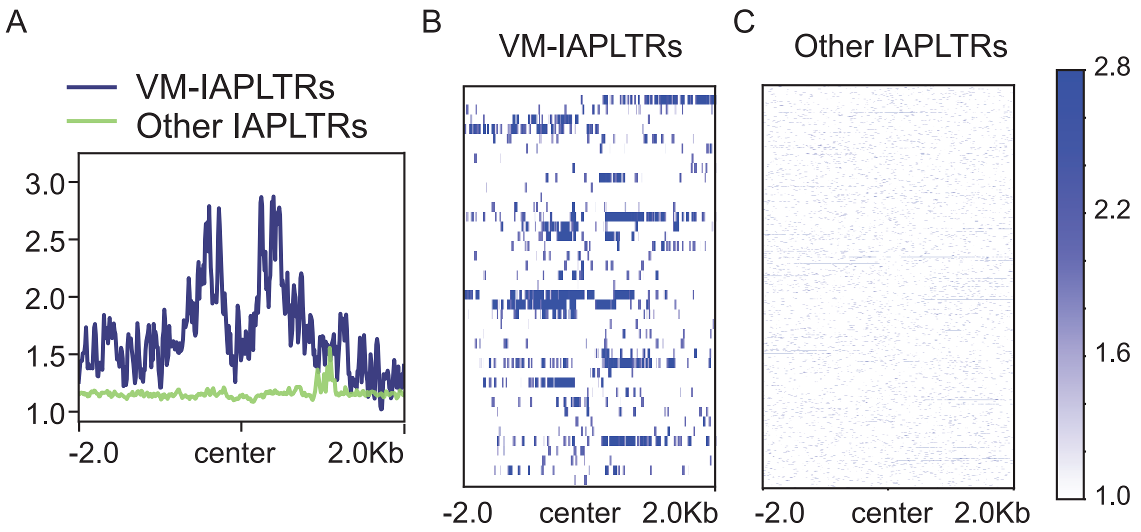

### Figure 2 - figure supplement 3

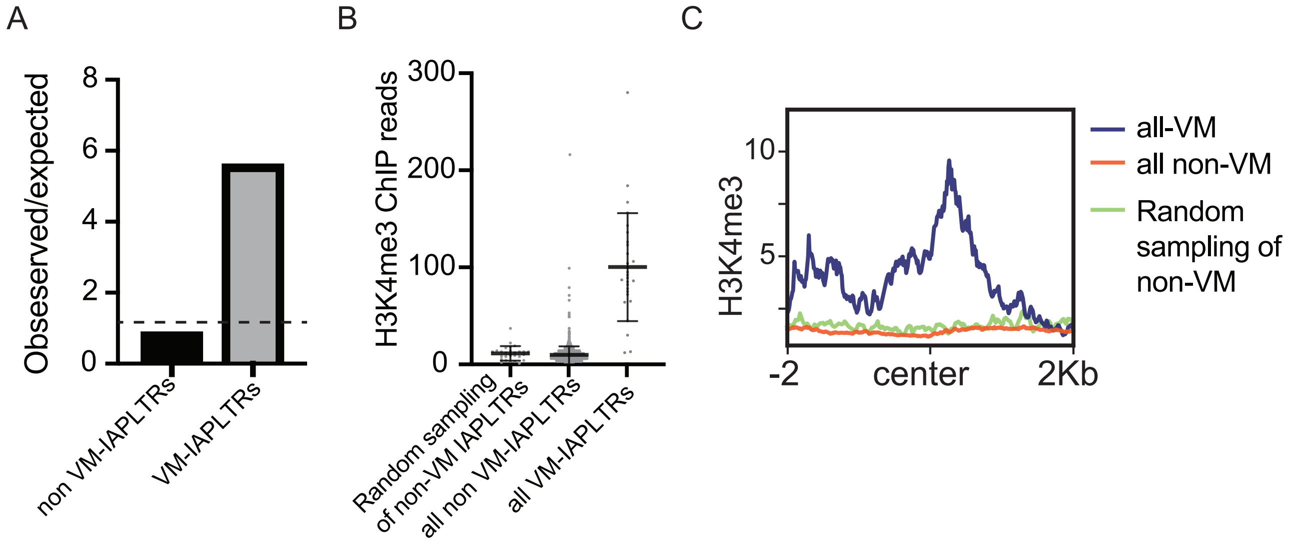

### Figure 2 - figure supplement 4

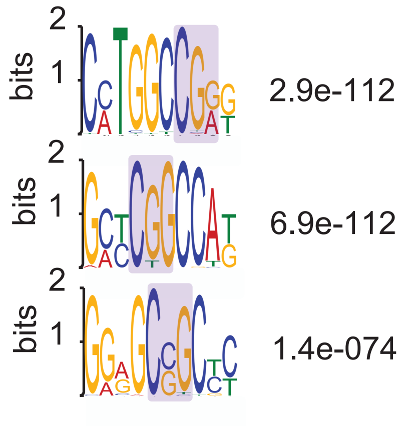

### Figure 3 - figure supplement 1

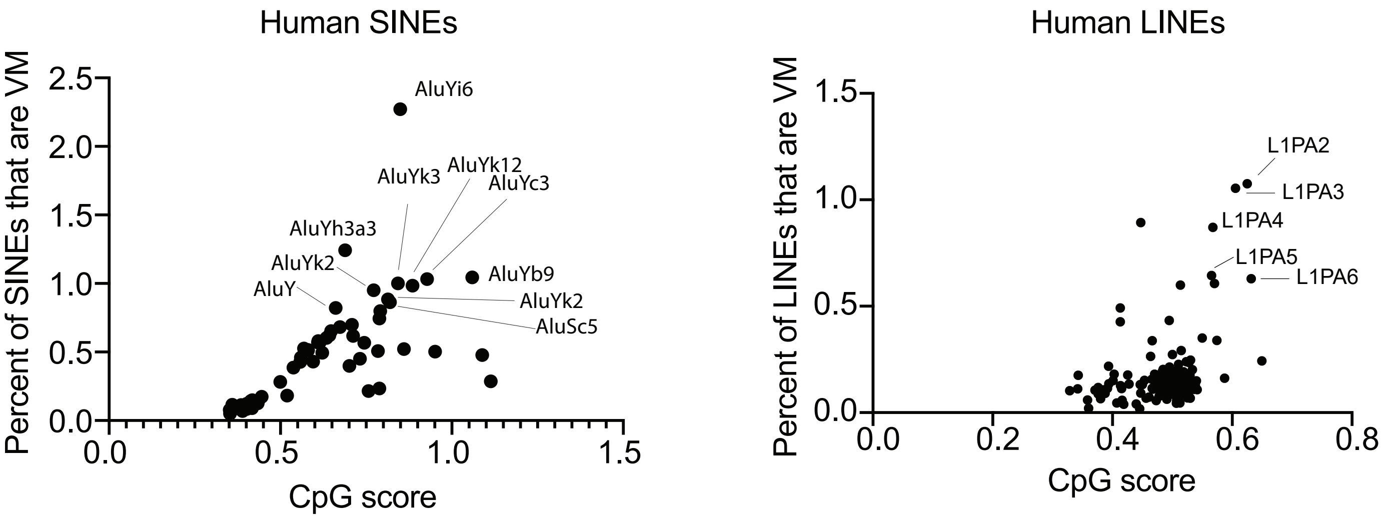

### Figure 4 - figure supplement 1

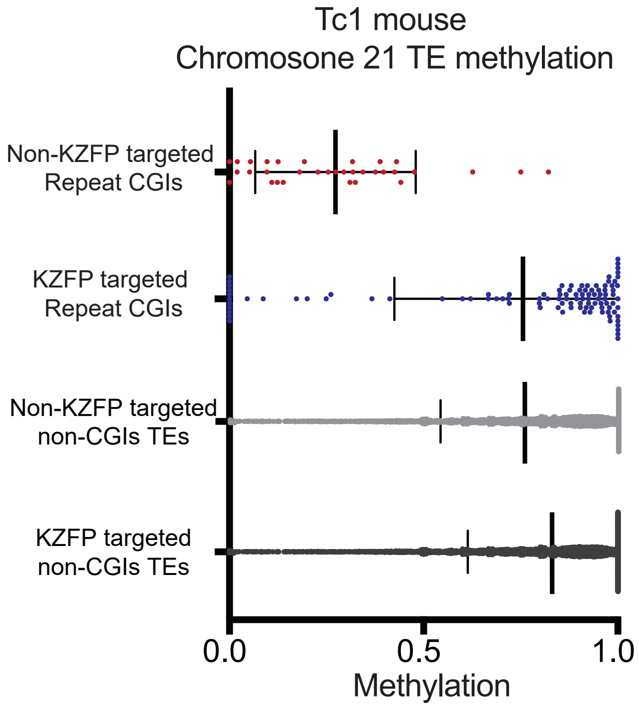

### Figure 4 - figure supplement 2

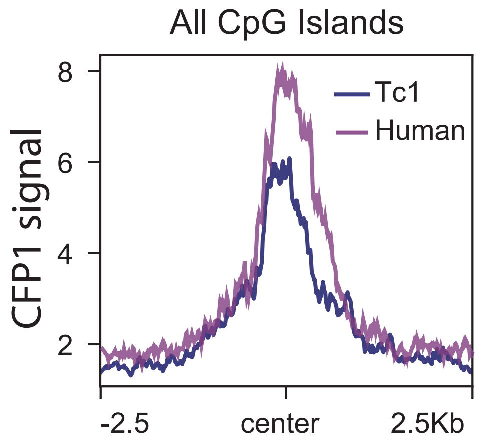

### Figure 5 - figure supplement 1

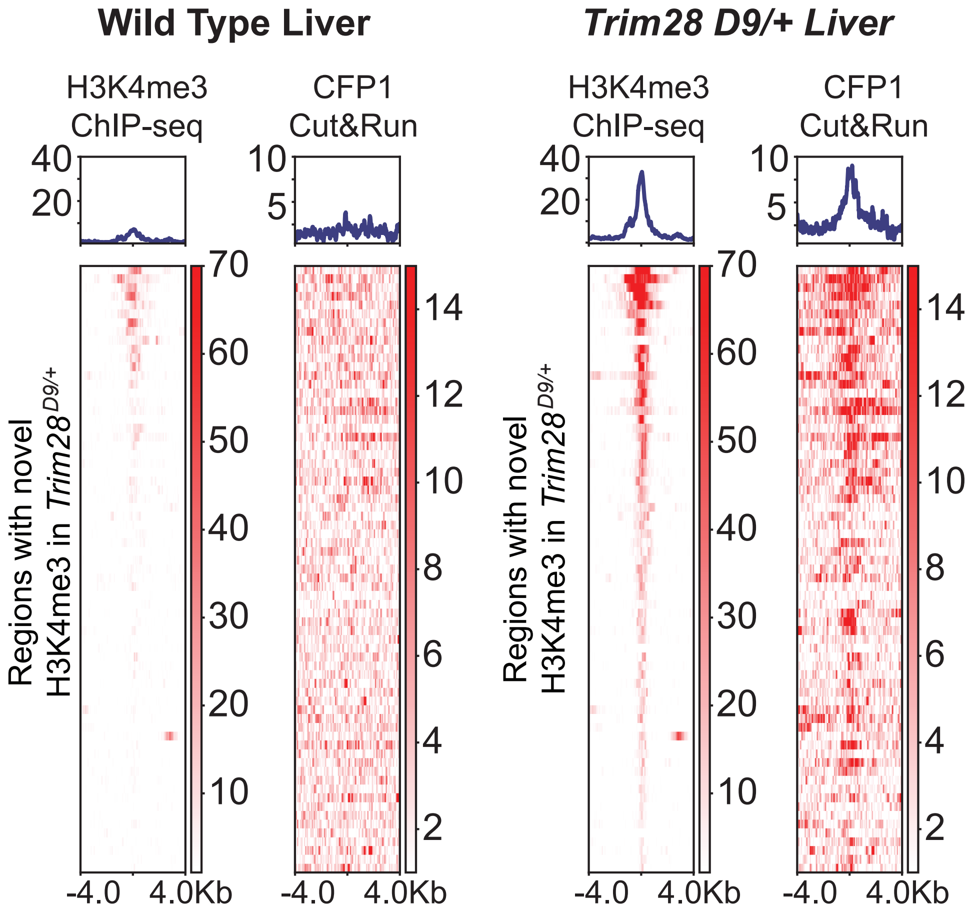

### Figure 5 - figure supplement 2

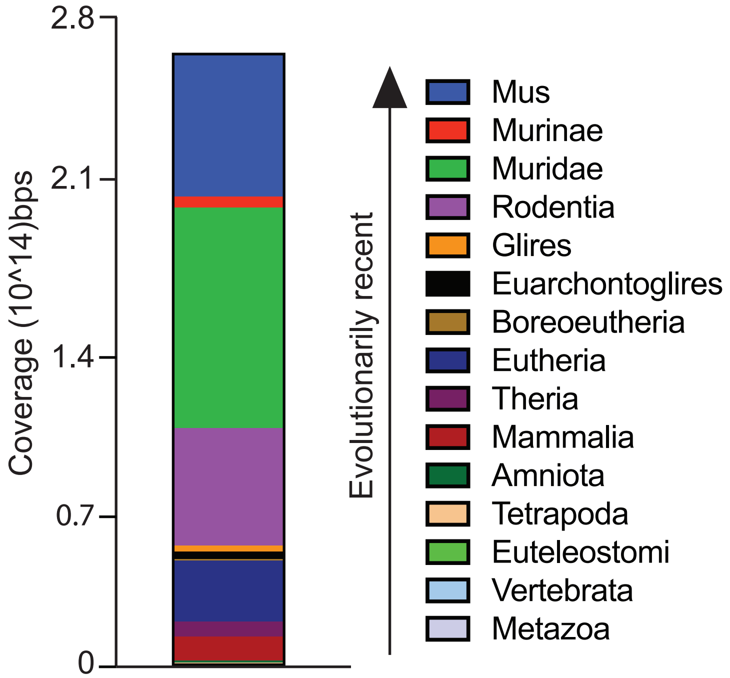

### Figure 6 - figure supplement 1

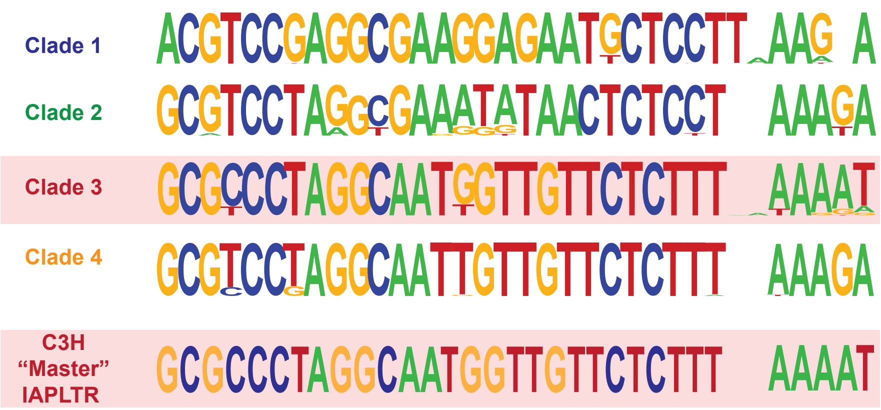
